# Regulatory effects of miRNA-19a on MAD2 expression and tumorigenesis in gastric cancer

**DOI:** 10.1101/2022.10.31.514416

**Authors:** J. Bargiela-Iparraguirre, J.M. Herrero, N. Pajuelo-Lozano, M. Perez, C. Cales, R. Perona, A.G. Quiroga, I. Sanchez-Perez

**Author notes:** Authors who equally contributed to this work (J. Bargiela-Iparraguirre, J.M. Herrero, N. Pajuelo-Lozano).

## Abstract

MAD2 is a key mitotic checkpoint protein that when overexpressed provokes chromosomal instability in gastric cancer. In this work, we used *in silico* analysis in combination with *in vitro* studies and clinical data to explore if miRNAs can regulate MAD2 at post-transcriptional level. By *in silico* analysis, we discriminate the expression of miRNAs between tumor and normal tissue, finding miR-19a and miR-203 targeted to 3’UTR *MAD2L1*. Luciferase Assays proved that those miR’s are specific to *MAD2L1* in human cells. RT-qPCR showed an inverse correlation between the expression miRNA19 and 203 and *MAD2L1* in a panel of gastric cancer cell lines and in a pilot series of patients’ study. The miR-19a expression reduces the migration ability of AGS cells and invasion in MKN45 cells. Furthermore, the expression of the miRNA in combination with mitotic checkpoint drugs increase apoptosis. Finally, the TCGA analysis showed that Gastric Cancer patients with overexpression of MAD2, showed higher overall survival when miR-19a was overexpressed. Together, our results defined miR-19a as a critical regulator of MAD2 protein in Gastric Cancer and could potentially be used as a prognostic biomarker in clinical use.

## 1. Introduction

Gastric cancer (GC) is the worldwide sixth most diagnosed cancer and the third leading cause of cancer deaths [1]. The treatment for this disease in its early stages surgery and the administration of adjuvant radio-chemotherapy. However, GC has a poor and late prognosis, mainly because of two reasons, the development of resistance after treatment and the lack of prognostic markers [2]. Different biomarkers, such as carbohydrate antigen CA-72-4, alpha-fetoprotein, carbohydrate antigen CA12-5, SLE, BCA-225, hCG and pepsinogen I/II, carcinoembryonic antigen (CEA) and CA19-9, are used in clinical practice for GC diagnosis, but none of those are efficient, highlighting the need for extensive research in this field [3].

Chromosome Instability (CIN) contributes to tumorigenesis because ends either gaining oncogenes or losing tumor suppressor genes, such as TP53, E1 D1 Cyclins and CDK6. This instability is caused by defects in spindle assembly checkpoint (SAC). The SAC is a protein complex formed by MAD1, MAD2, BUB1, BUBR1 and BUB3 proteins which prevents anaphase until all chromosomes are attached to a bipolar mitotic spindle[4]. MAD2 is one of the key proteins in the SAC complex [5]. Despite the controversy in the literature regarding the MAD2 function in tumor prognosis, there is a growing body of data indicating that MAD2 plays a role in cancer disease, for example the high expression of MAD2 is directly related to the occurrence of aneuploidy and tumorigenesis [6, 7]. In this line, our research group demonstrated that overexpression of MAD2 correlates with tumorigenesis in GC. Furthermore, interference of *MAD2L1* by using sh*MAD2L1*, decrease migration and tumor growth. [8, 9]. These data encouraged us to study the regulation of MAD2 levels due to posttranscriptional regulation by MicroRNAs (miRNAs).

MicroRNAs (miRNAs) are endogenous, small non-coding RNAs that function in regulation of gene expression through the binding to a canonical sequence 7-8 nucleotides in the 3’UTR end of the mRNA, preventing translation of the protein. In cancer, the altered expression of miRNAs modifies the regulation of oncogenes and / or tumor suppressor genes [10-12]. The public bioinformatics databases like GEO (Gene Expression Omnibus) and TCGA (The Cancer Genome Atlas) can be used to screen significant genetic or epigenetic variations occurring in carcinogenesis [13]. miRNAs profiling and deep sequencing provided direct evidence that miRNA expression is dysregulated in cancer and its signatures could be used for tumor classification, diagnosis, and prognosis [14]. The modulatory roles of miRNAs in GC carcinogenesis have been also reviewed [15]. For instance, downregulation ofmiR-203 and miR-214 expression in cancer tissues compared to healthy [16, 17] are related with CHK1 overexpression and radiotherapy resistance in GC [18].

In this work, our hypothesis is that MAD2 overexpression in GC might be caused by miRNA deregulation. Our starting point was the bioinformatic and functional analysis which provided miR-19a and miR-203 as candidates to downregulate *MAD2L1*. Our experiments indicates that miR-19a expression decreases tumor cell migration and invasion, and synergized cell death after treatment with different drugs. In addition, the analysis in a pilot series of patients demonstrated the inverse correlation of miR-19a and MAD2 protein expression. Finally, the Kaplan-Meier analysis showed that patients with high miRNA-19a levels had a longer overall survival (OS) than those who had low miRNA-19a levels. Therefore, we suggest that miRNA-19a could be considered in preclinical studies as a biomarker for prognosis and therapy response.

## Materials and Methods

### Gene expression profile analysis

The miRNA expression data in a large set of GC patient samples as down-loaded from the GEO (http://www.ncbi.nlm.nih.gov/geo/) and analyzed using custom R scripts for statistical programming http://www.r-project.org. Briefly, we compared probe values in each sample group (“normal”, GC) using a Student’s t-test and the resulting p-values were adjusted for multiple testing by the Bonferroni method.

### RT-qPCR

Total RNA was extracted from cells and tissues using Trizol Reagent (Invitrogen, Carlsbad, CA) according to the manufacturer’s protocol. Mature miR-19a, miR-203 and miR-224 were quantified by TaqMan MicroRNA Assays, according to manufacturer’s instructions (REF: hsa-miR-19 000395, hsa-miR-203 000507 and hsa-miR-224 000599). Mature miRNA expression was normalized to snRNA-U6 (REF: 001973). The relative changes in gene expression were analyzed by the 2−ΔΔCT method. *MAD2L1* was detected by RT-qPCR analysis using the SYBR Green detection system (Roche Applied Science) and normalized to the level of the GAPDH mRNA expression and using the 2-ΔΔCT cycle.

### Clinical tissues samples

A total of 44 pairs of human GC tissues matched with adjacent non-cancerous gastric tissues were obtained from patients in INCAN (Instituto de Cancerología). The RNA was donated by INCAN for which we acknowledge.

### Cell cultures, transfections and treatments

Five GC cell lines, AGS, HGC27, Hs749t, MKN45 and ST2957 and 293T were grown in RPMI-1640 or DMEM supplemented with 10%(vol/vol) fetal bovine serum (FBS). Cultures were maintained at 37ºC in a humidified atmosphere with 5% CO_2_. Cells were seeded in 12 well plates and transfected using Lipofectamine 2000 (Invitrogen) according to the manufacturer’s instructions. CDDP was donated from Ferrer FARMA. Paclitaxel was acquired from Sigma-Aldrich. **I5** (*trans*-[PtI_2_(ipa)_2_]) & **I6** (*cis*-[PtI_2_(ipa)_2_]), where ipa stands for isopropylamine, were synthesized and characterized as indicated in the references [19, 20].

### Luciferase reporter assay

The 3’ UTR of the human *MAD2L1* with predicted miR-binding site was amplified from HCT116 genomic DNA and cloned to the psiCHEK2 located downstream of the Renilla translational stop codon into XhoI/NotI restriction enzymes (Promega). 3’UTR mutant was obtained by using the Site Directed Mutagenesis (New England Biolabs). miR-19a, miR-203 and miR-139 were cloned according to manufacturer’s instructions in the miRNASelect™ pEGP-miR vector. pEGP-Null was used as a control (Cell Biolabs. INC, San Diego, USA). Firefly and Renilla luciferase activities were measured 48h after transfection using the Dual-Luciferase Reporter Assay Kit (Promega), according to the manufacturer’s instructions. Luciferase units were register in a GloMax® 96 Microplate Luminometer.

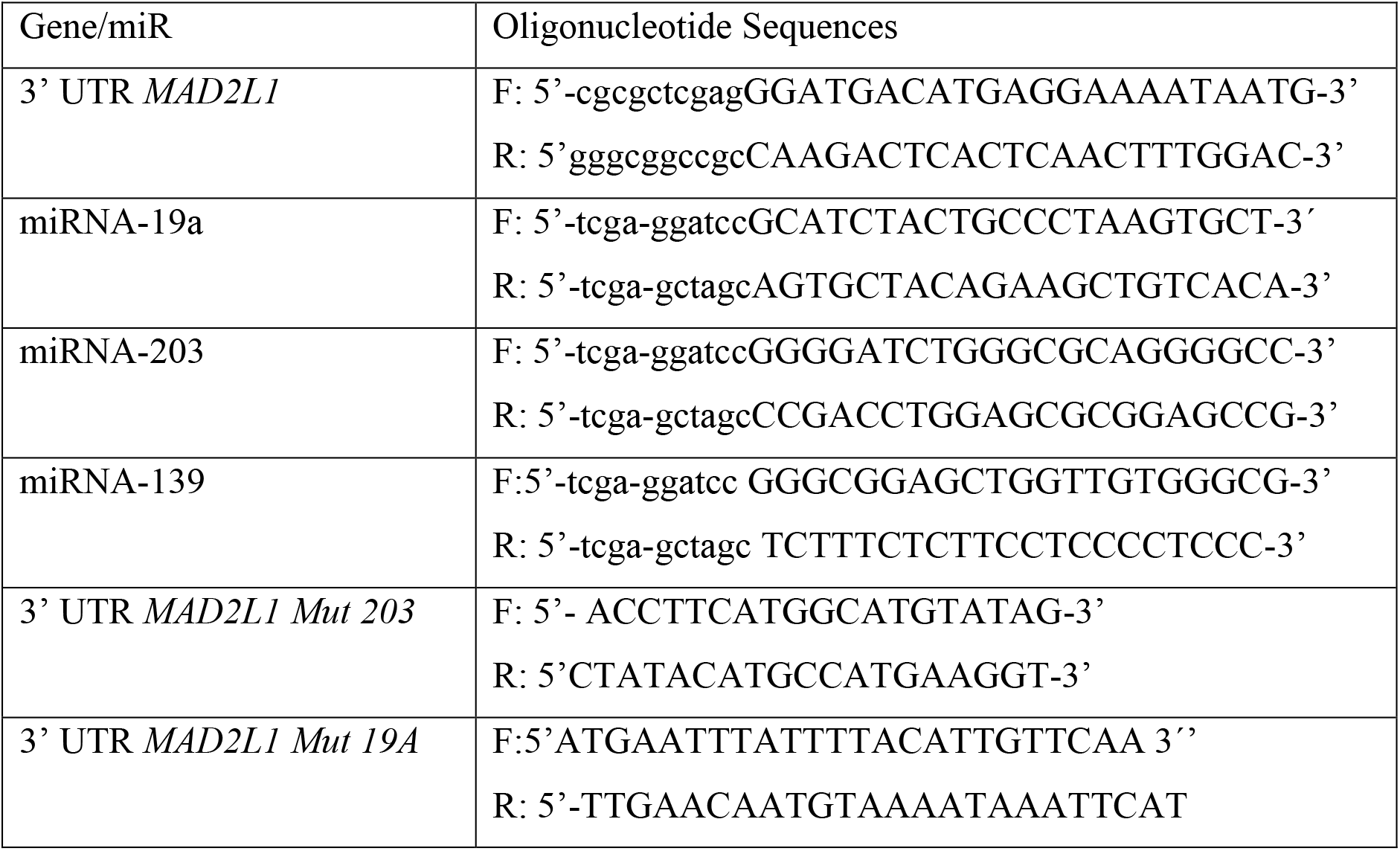

### Cell Cycle profile analysis

Cells were fixed with 70% cold EtOH and stored at 4 ºC overnight. Cells were incubated with Propidium Iodide (50 μg/mL) and RNAse (10 μg/mL) for 30 min and DNA content was evaluated using FACS CANTOII Flow Cytometer (Becton Dickinson). Results were analyzed using FACS Diva Software. IC_50_ was used as a dose for drugs treatments based on our results for these cell lines. Cisplatin: AGS 20μM; MKN45 10μM. I5: AGS 40μM; MKN45 30μM. I6: AGS 50μM; MKN45 50μM.

### Western Blotting

Total protein extracts were obtained using the previously described lysis buffer [21].Twenty μg of protein extracts per sample were loaded in 15% SDS-PAGE polyacrylamide gels, and then transferred onto PVDF membranes, followed by immunodetection using appropriate antibodies. Antibodies against the following proteins were used: MAD2 (1:1000, sc-28261) was purchased from Santa Cruz Biotech-nology; and α-Tubulin (1:10000, T9026) was obtained from Sigma. Horseredish perox-idase-conjugated secondary antibodies anti-Mouse IgG (1:5000) were purchased from Santa Cruz Biotechnology, and anti-Rabbit IgG (1:5000) from Bio-Rad Laboratories.

### Cell migration and invasion assays

For the wound-healing assay, a confluent monolayer of AGS cells was scratched into a 24-well plate with a sterile tip. The cell migration’s ability to fill the wound was studied up to 40h. The relative distance traveled by the leading cell edge was assessed by time-lapse microscopy using the Cell Observer Z1 (Zeiss) at 37ºC and 5% CO_2_/95% air, using the imaging software Axiovision 4.8 and the Cascade 1 k camera. Images were taken every 2h and further processed using the Digital Image Processing Software AXIOVISION (Zeiss). The wound closure was quantified by measuring the remaining unmigrated area using ImageJ. Invasion assays were performed in a cell culture insert with transparent PET membrane (6.5 mm diameter, 8.0 μm pore size, transwell migration and invasion assays (Falcon, Corning Inc., Corning, NY, USA)), coated with 6 μg growth factor reduced Matrigel (BD Biosciences, Bedford, MA, USA). MKN45 cells (1.5×10^5^) were seeded in the upper chamber. FBS (20%) was used as a chemoattractant and placed in the bottom chamber. Cells were allowed to invade for 48h at 37°C with 5% CO_2_. Then, non-invading cells were removed using a cotton swab, and the filters were stained with Diff Quik (Dade Behring, Newark, DE, USA). Invasive cells were counted in 50 fields of maximum invasion under a light microscope at 40× magnification. Representative images were captured using an Axiophot Zeiss microscope.

### Statistical analysis

Statistical analyses were performed using Graphpad prism 5.0 or IBM SPSS 22 software, ANOVA or Student’s 2-tailed t-test. Values of *P < 0.05 were considered significant.

## Results

### miR-19a and miR-203 modulates MAD2L1 levels in GC

We decided to investigate the expression of miRNAs in samples from human GC to find miRNA candidates that could regulate MAD2 expression by using the public dataset GSE30070, from GEO database. This dataset describes the miRNA expression profile in 90 GC samples, collected prior chemotherapy treatment, and 34 samples from healthy volunteers. The diagram overview of the analysis is depicted in Figure 1A. The study gave us 97 hits differentially expressed between normal and tumor samples (adjusted p-value < 0.05). These hits were grouped according to the corresponding miR ID annotation in the GLP13742 platform (Agilent-015868 Human miRNA Microarray (miRBase release 9.0 Feature Number version)). The hits corresponded to 29 hsa-miRNA (Table SMI). Then, Target SCAN & mirDB (MicroRNA Target Prediction Database) tools predicted that from the 29 miRNA ID only few of them showed *MAD2L1* as a possible target (Table SM2). Figure 1B shows the interaction site with *MAD2L1* 3’UTR for some miRNA, the specific details of miR-19a, miR-203, and miR-224 and their statistical data obtained from the *in silico* study.

**Figure 1.**
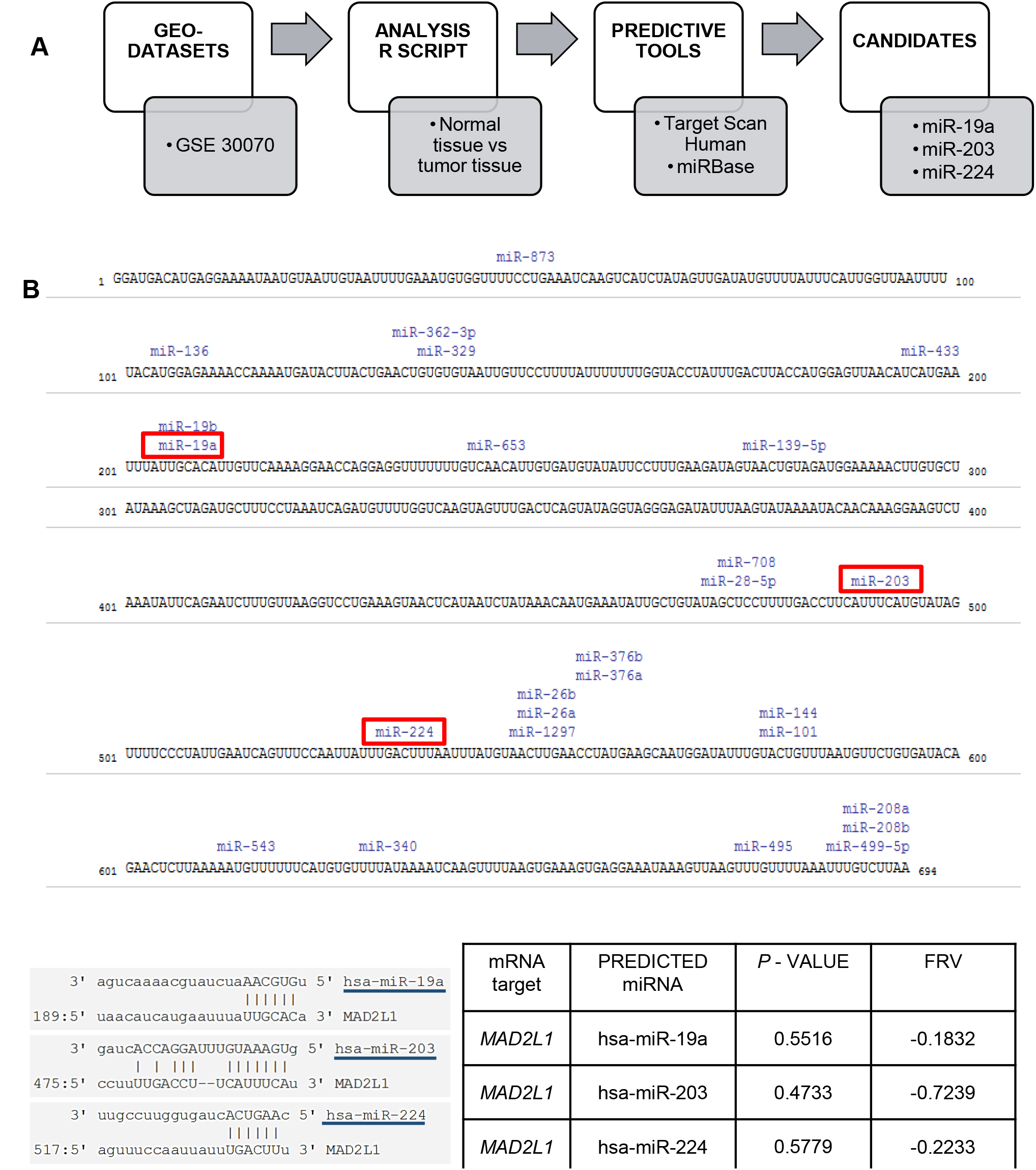
miRNAs selected candidates for *MAD2L1* regulation. **A)**. Schematic Diagram of *in-silico* analysis. Public GSE dataset (GSE30070) was selected for differential expression analysis, and those miRNAs showing significant differences between groups were selected. The miRNAs with best p-value were analyzed by predictive tools of different databases to predict MAD2-targeting miRNAs. Best candidates were miR-19a, miR-203 and miR-224 binding sites on human 3’UTR *MAD2L1* mRNA. Graph shows the 3’UTR*MADL1* sequence and the miRs binding sites (the selected miRs were reflected with red boxes).

Next, we proved their potential action as posttranscriptional regulators of *MAD2L1* by using the luciferase reporter assay. 293T cells were transfected with psiCHECK-3’UTR *MAD2L1* and pEGP-miR-19a or pEGP-miR-203. We observed a reduction of luciferase activity upon expression of miR-19a and miR-203 (Fig. 2A, left panel) whereas the co-transfection with psiCHECK-3’UTR *MAD2L1* mutant for miR-19a or miR-203 binding sites did not affect the luciferase activity (Fig. 2A, right panel), hinting that miR-203 and miR-19a bind to 3’UTR site of *MAD2L1*. In addition, we included the expression of mir-139-5p, because its interaction with 3’-UTR *MAD2L1* has been described[22]. But in our experiment, we observed no effect on luciferase activity after expression of miR-139, indicating that no interaction occurs on 3’UTR-*MAD2L1*. (Fig. SM1A,B)

**Figure 2.**
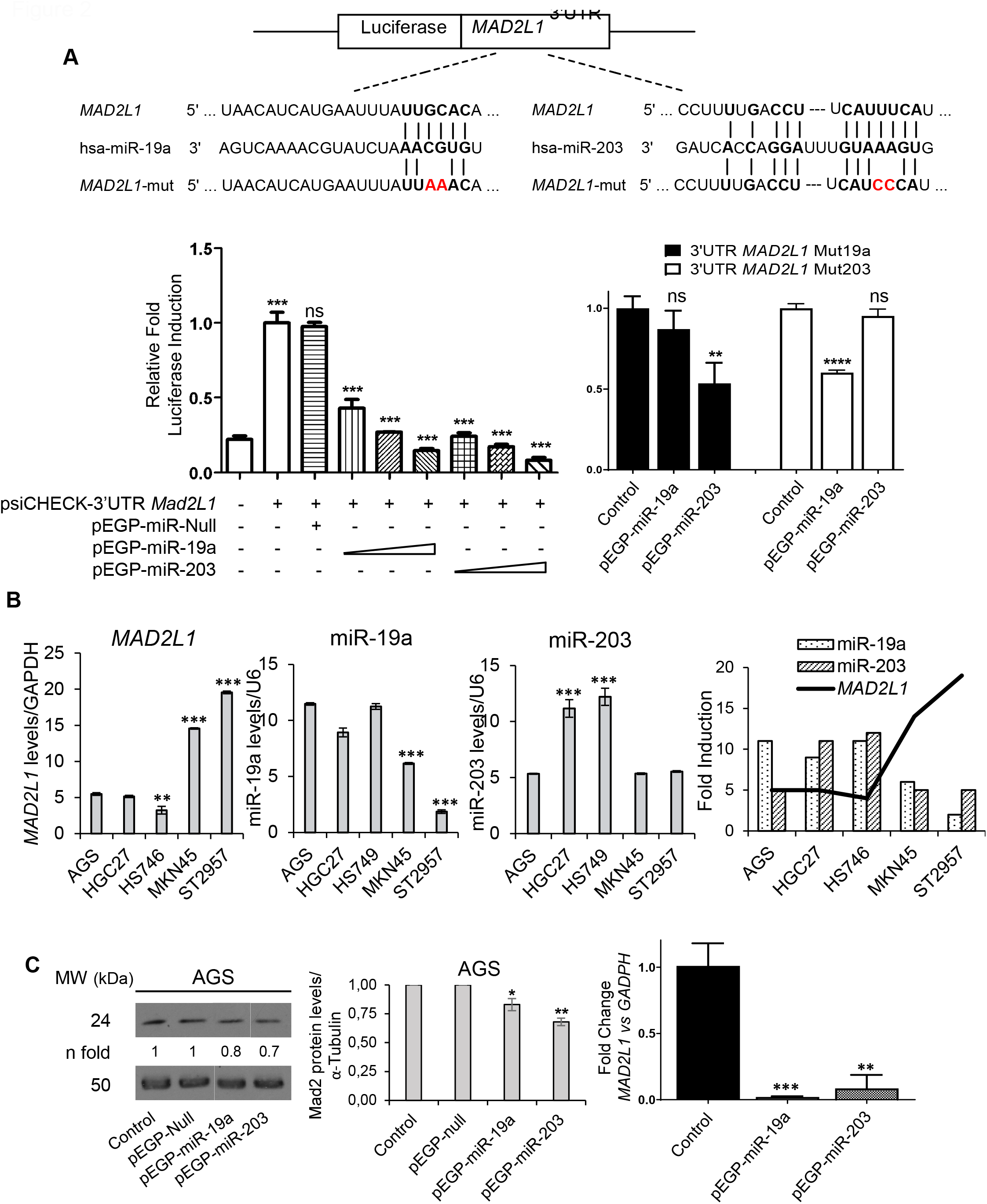
miR-19a and miR-203 interact with 3’ UTR-*MAD2L1*. **A)**. Schematic Diagram of Luciferase assay in: (Left Graph) 293T cells co-transfected with pEGP-Null or different doses of pEGP-miR-19a and/or pEGP-miR-203 and psiCHECK-3’UTR *MAD2L1* reporter; (Right Graph) Luciferase assay in 293T cells co-transfected with pEGP-Null or pEGP-miR-19a or pEGP-miR-203 and psiCHECK-3’UTR *MAD2L1*mut19a (■) or psiCHECK-3’UTR *MAD2L1*mut203 (□) reporter. In both experiments luciferase activity was represented as a fold induction compared to untransfected cells. The experiments were performed three times, and statistical differences were assessed by one-way ANOVA (*P<0.05, **P<0.005, ***P< 0.001). **B)** RNA was isolated from AGS, HGC27, HS749, MKN45 and ST2957 cell lines. *MAD2L1*, miR-19a and miR-203 were quantified by RT-qPCR. *MAD2L1* was normalized with GAPDH and miRNAs with U6, respectively. The statistical significance was evaluated with one-way ANOVA (**P<0.05 and ***P<0.001). The experiment was performed three times. In the right side we reflect a comparison between *MAD2L1* levels and both miRNAs in the studied GC cells. **C)** MAD2 expression normalized with α-Tubuline *via* Western Blotting (left) and *MAD2L1/GADPH* fold-change by RT-qPCR (right) in AGS cells without and after transfection with both miRNAs and an empty vector as control. The statistical significance was evaluated with Student’s 2-tailed t-test (*P<0.05, ** P<0.005 and ***P<0.0005)

We wanted to know the ratio of *MAD2L1* and miRNA expression in GC. To that end, we evaluated by qPCR the expression levels of *MAD2L1*, miR-19a, miR-203 and miR-224 in AGS, HGC27, Hs746t, MKN45 and ST2957 cell lines. According with our previous data, MKN45 and ST2957 showed higher expression of *MAD2L1* [15] than AGS, HGC27 and HS749 cells. By contrast, expression of miR-19a or miR-203 were lower in MKN45 and ST2957 than in the other cell lines (Fig. 2B). These data suggest an inverse correlation between *MAD2L1* expression and miR-19a or miR-203 expression which we indicate with a bold line in Figure 2B (right graph). No correlation exists with miR-224 (Fig. SM1C) and as a result miR-224 was discarded for the rest of the experiments. Furthermore, we corroborate the role of miR-19a or miR-203 in AGS cells. Western Blot analysis showed a reduction on MAD2 protein when miRs are transfected. Same results were observed studying the *MAD2L1* gene expression (Fig. 2C), however expression of miR-139a do not have affect over MAD2 protein expression. Based on such expression and the previous luciferase assay, we can discard miR-139 for the rest of the experiments and focus on miR-19a and miR-203. The results support the hypothesis that both miRs contribute to the regulation of MAD2 protein levels in GC cells.

### miR-19a expression modulates apoptosis of GC Cells treated with chemotherapeutic agents

MAD2 is key for mitosis control, so we studied the effect of miRNA expression on the cell cycle profile. We observed that apoptosis is increased in AGS and MKN45 cells, after expressing miR-19a or miR-203 (Fig. 3A) maybe due to a leak mitotic checkpoint.

**Figure 3.**
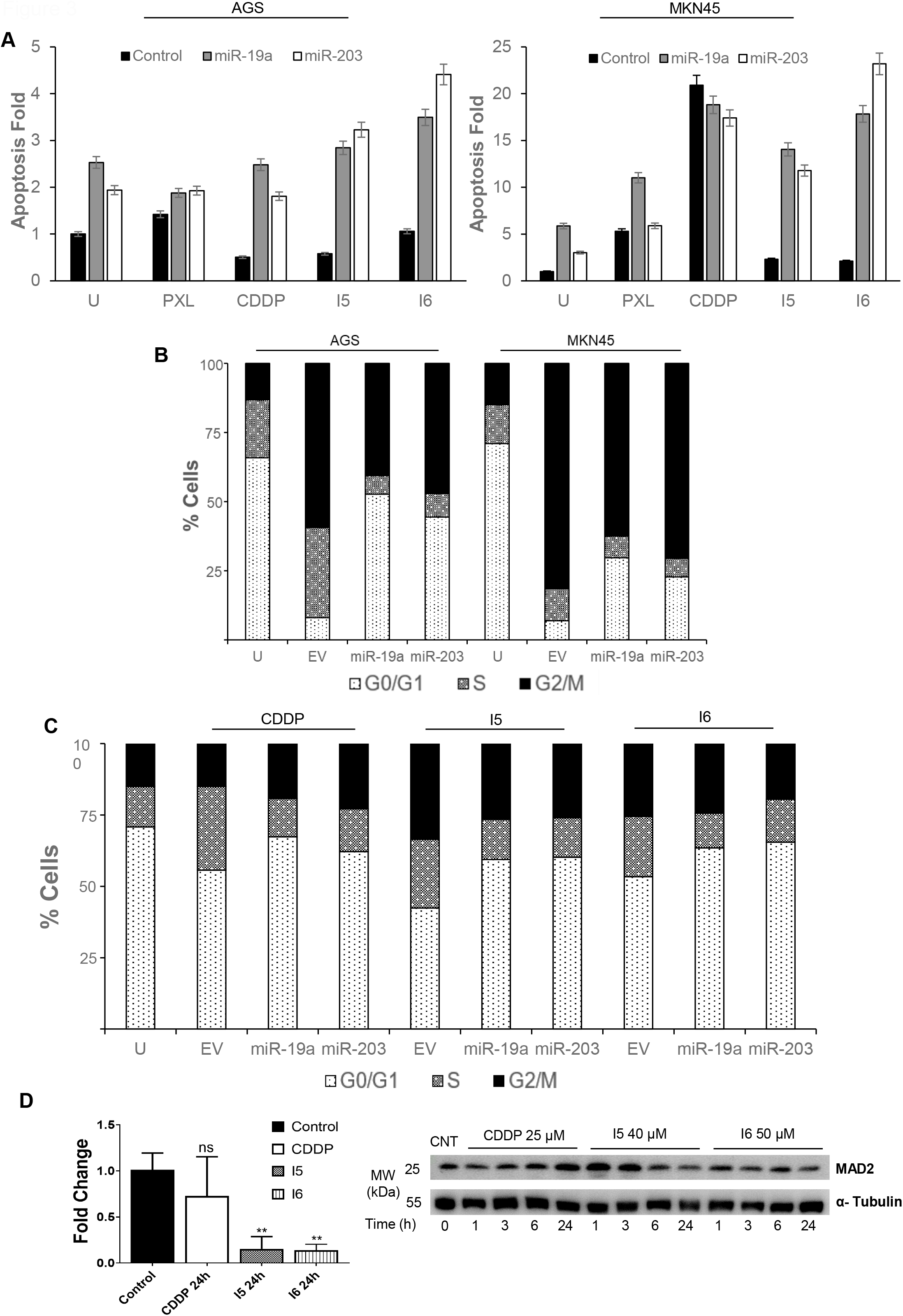
mi-RNAs modulates apoptosis and cell cycle profile. **A)** AGS and MKN45 cells were transfected with pEGP-null (EV), pEGP-miR-19a or pEGP-miR-203. 24h after transfection, cells were treated with the drugs (0.1 μM PXL, 10 μM CDDP, 30 μM I5 and 50 μM I6) for 24h, harvested and fixed with EtOH/PBS (see M&M for details). Graph shows the fold induction of apoptosis referred to control untreated cells. **B)** AGS and MKN45 cells were transfected as in A). The graph shows the percentage of cells in each phase G1, S and G2/M after PXL treatment (0.1 μM). **C)** MKN45 cells were transfected as mentioned above. Picture shows the cell cycle profile after treatment with CDDP (10 μM), I5 (30 μM) and I6 (50 μM). DNA content was assessed by flow cytometry, and cell cycle distribution was analyzed using FACS Diva software. All the experiments were performed three times and it is represented the mean values of these three days. D) *MAD2L1/GADPH* fold-change was analyzed using RT-qPCR (left) and MAD2 expression normalized with α-Tubuline by Western Blotting (right) in AGS cells without and after treatment with the IC50 of CDDP, I5 & I6 at different times (1, 3, 6 and 24h in Western Blot, and 24h in PCR). The statistical significance was evaluated with Student’s 2-tailed t-test (*P<0.05 and ** P<0.005)

We wonder if miRNA could enhanced the apoptosis induction after treatment with clinical drugs (PXL, microtubules polymerization inhibitor arresting the cell cycle in mitosis and CDDP, covalent drug that affect mainly S-phase), as well as with two cisplatin derivatives with iodide ligands (I5 & I6, which have been reported a G2/M arrest [20]). Importantly, the treatment with PXL, CDDP, I5 or I6 potentiates the cell death, being synergistic in cells treated with the new platin-derived agents I5 and I6 (Fig. 3A). Regarding the effect of the miRNAs expression on the G2/M cell cycle phase after drugs exposure, it was observed that in the presence of PXL, miRNA expression reduced the percentage of G2/M cells in both cell lines (Fig. 3B). Same trend was observed with the platinum iodido complexes (I5 & I6) in the MKN45 cells (Fig. 3C). We observed that I5 and I6 drugs decrease the transcription of *MAD2L1* and MAD2 protein expression in AGS cells (Fig. 3D). This data indicate that I5 & I6 compounds with the miR expression could synergize the downregulation of MAD2, and its effect on mitotic checkpoint..

### miR-19a modulates tumorigenesis of GC Cells

Next, we studied the role of these miRNAs (19a, 203) on GC migration and invasion by using wound healing analysis and Transwell assay, respectively. We expressed the miR-19a, miR-203 in the AGS cell line. After 40h, our analysis showed that miR-19a expression reaches 40% Wound Closure compared to the complete closure in control cells. However, miR-203 expression had few effects on migration (Fig. 4A). To further confirm a general role of miR-19a in GC tumorigenesis, we analyze the invasion capability in MKN45 cells. We observed a reduction in the average value for invasion in cells expressing both miRNAs (Fig. 4B); those values change from 42 in control cells to 28.5 with miR-19a and to 29 in miR-203 expressing cells.

**Figure 4.**
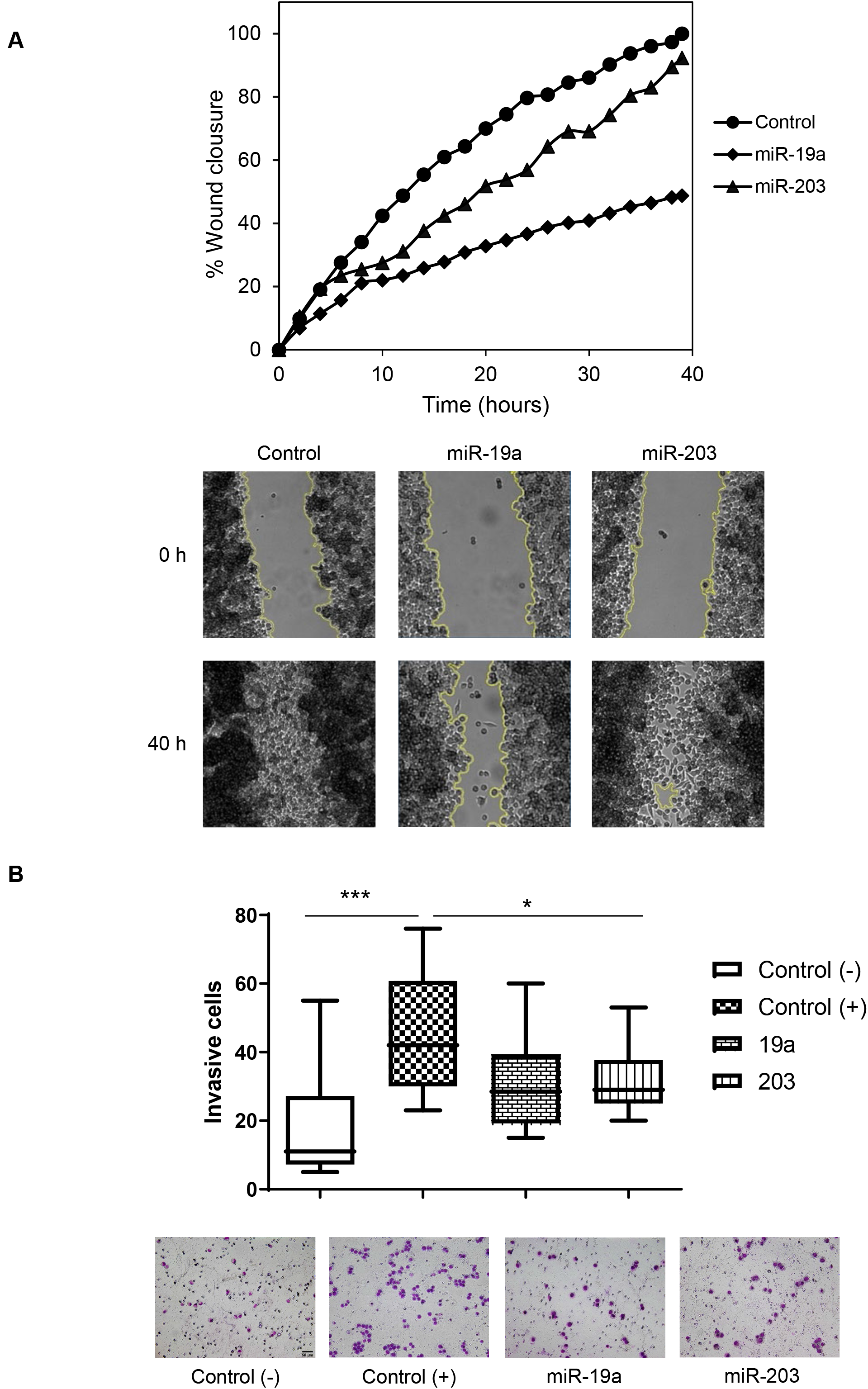
miR-19a reduces migration and invasion in GC cells. **A)** AGS cells were transfected with miR-19a and miR-203. The graph shows the percentage of wound closure over the study time using the ImageJ program. AGS (●), miR-19a (♦) and miR-203 (▲). The experiment was performed twice, and statistical differences were assessed by one-way ANOVA (*P < 0.05, ** P < 0.005, *** P <0.001). Representative images of the first (0h) and last picture (40h) of control AGS, pEGP-miR-19a and pEGP-miR-203 transfected cells taken during the wound healing experiment. Images were taken at 10× magnification, every 2h for 48h. The yellow line represents the wound border. **B)** MKN45 cells were transfected as in A) The graph shows the quantification of stained invasive cells from MKN45 and MKN45 miR-19a/203 using the transwell assay, without and with 20% FBS at 48h. Representative photographs of the experiment are shown on the right. Scale bar: 50 μm. Statistical differences were assessed by one-way ANOVA (*P < 0.05).

These results indicate that the miR-19a expression might regulate Mitotic Checkpoint (MC) and tumorigenesis, mainly due to MAD2 downregulation in GC cells.

### Relevance of MAD2 Expression in Gastric tumors

To confirm the *in cellulo* results we evaluated the expression and the clinicopathological characteristics of GC patients from a cohort of 44 patients divided into two groups of population: high *MAD2L1* (n=17) and low/normal *MAD2L1* expression (n=27). The general features of the 44 patients are summarized in Table 1. No significant association between *MAD2L1* levels and age, sex, stage, Lauren and Her2+ expression or PFS (progression free survival) was observed in patients with GC (p< 0,05). However, we observed significant differences in mortality, also reflected on the Kaplan-Meier curve, that showed early differences in Overall Survival (OS) between both populations (Fig. 5A). The survival median of the patients with low *MAD2L1* reaches 54 months, while those with high *MAD2L1* median reaches 20 months. We also checked the OS in a wide panel of patients (n=592) available in KM plot (excluding GSE62254 dataset as the webpage suggested due to it has different characteristics), obtaining the same trend: the survival median of the patients with low *MAD2L1* reaches 29.5 months, while those with high *MAD2L1* median reaches 19.5 months (Fig. 5B). Furthermore, we studied the correlation between miR-19a and miR-203 and *MAD2L1* mRNA levels in a short subgroup of 9 GC patients from the 44 patients indicated above. We observed that GC Tissues of patients with high *MAD2L1* expression showed lower levels of miR-19a and miR-203 (Fig. 5C). These results suggested that the inverse correlation between miR-19a / miR-203 and *MAD2L1* expression detected in GC cells (Fig. 2B) is also evident in GC patients’ samples.

**Table 1.**
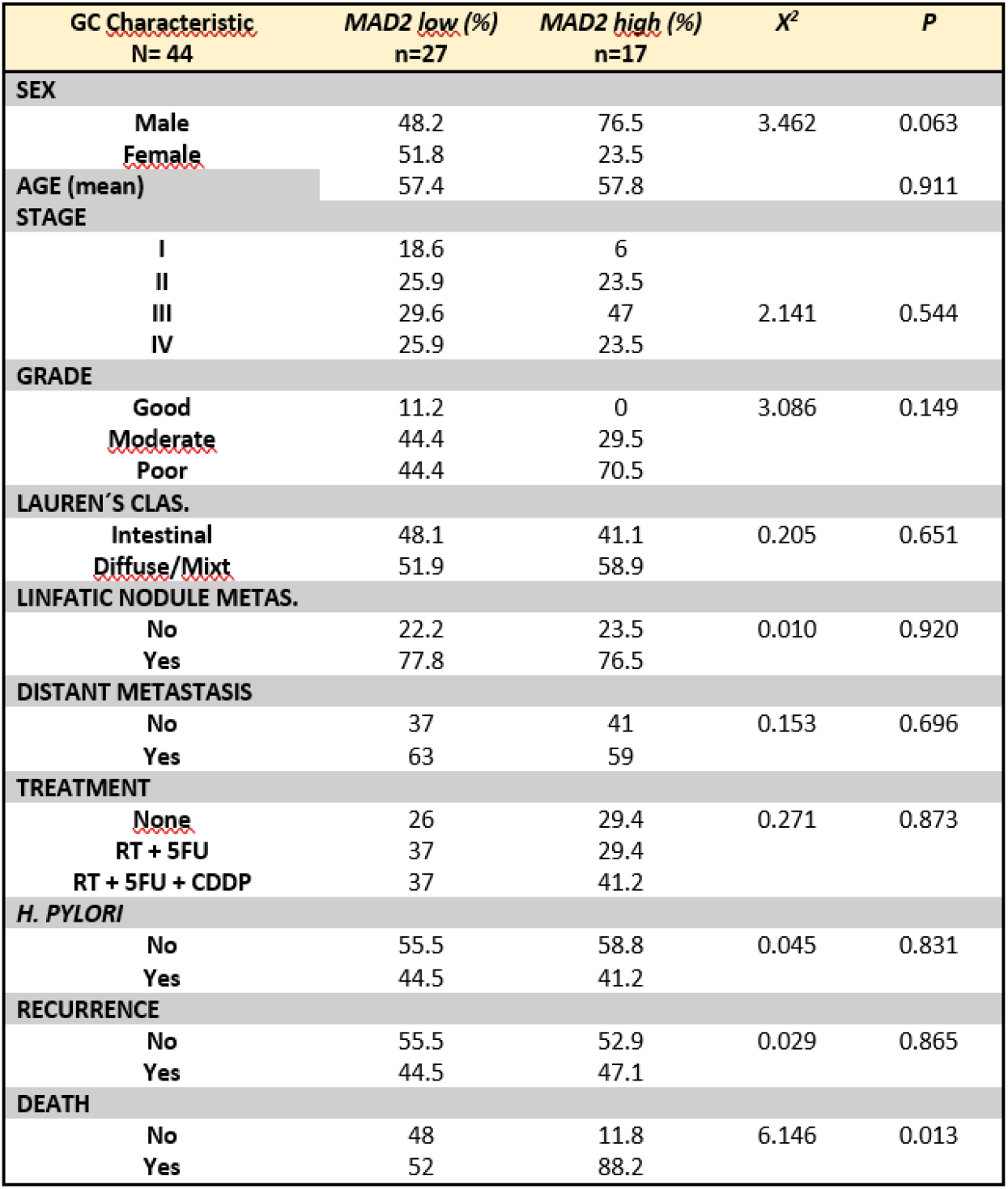
Association of *MAD2L1* expression with the clinical and pathological characteristics of the selected 44 GC patient

**Figure 5.**
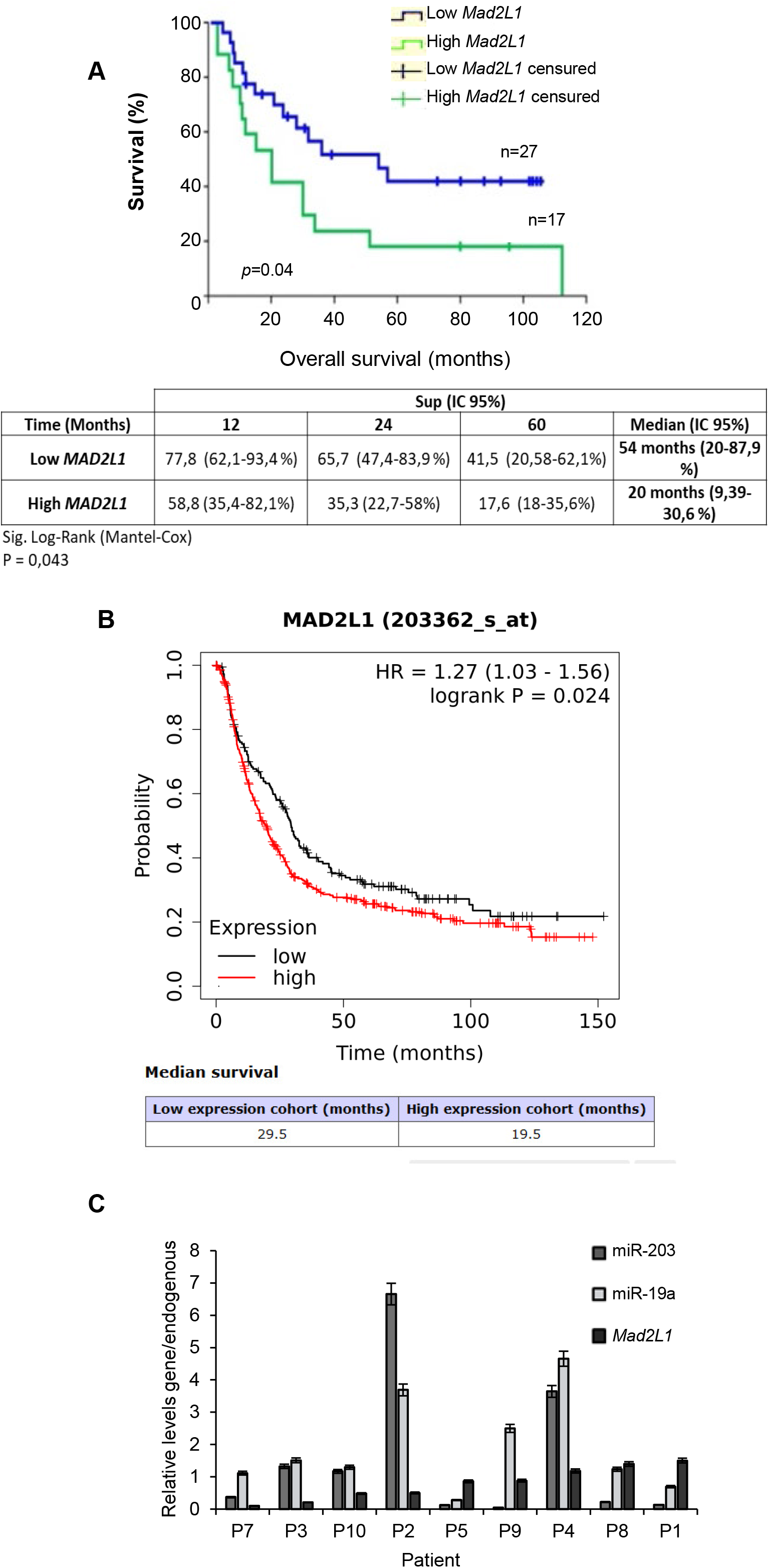
*MAD2L1* levels correlates with OS. **A)** Kaplan-Meier analysis of 10-years overall survival curves of 44 GC patients divided in high *MAD2L1* (n=17) versus low *MAD2L1* expression (n=27) with respect to expression level of MAD2 of GC patients. The statistical significance was evaluated with Mantel-Cox test. B) Kaplan-Meier analysis of 592 GC patients divided in high *MAD2L1* (n=406) versus low *MAD2L1* expression (n=186) generated in kmplot.com by using MAD2L1 (203362_s_at) as a target gene. **C)** RT-qPCR in nine tissue samples of GC of *MAD2L1*, miR-19a and miR-203; P<0.05 vs adjacent normal tissues samples.

### miR-19a as a predictive biomarker in Gastric tumors

Based on the above results, we aimed to study the potential used of miR-19a as a new GC biomarker. Firstly, we tested the association of miR-19a with the survival performance of patients with GC by using the log-rank test and Kaplan-Meier curve. The GC samples (TCGA: n=372) were divided into two different groups according to miR-19a expression. The mean value was used as cut-off value (p<0.05): high expression group (n=122) and low expression group (n=250). Based on Kaplan-Meier analysis, the OS rate of the low group was markedly worse than that of the high group, with a p value equal to 0,021 (Log Rank –Mantel-Cox), suggesting that down-regulation of miR-19a may contribute to the malignant progression of GC (Fig. 6A). In addition, we divided the dataset into two groups: high and low *MAD2L1* expression, according to ROC Curve. Then we assessed the predictive prognosis capacity of miRNA-19a, in high *MAD2L1* group. We found the OS for 1, 2 and 5 predictions years were, respectively 84.2%, 64.0%, 56.9%, in the high expression group, better than data found in the low expression group 73.9%, 48.6%, 30.5%. (Fig. 6B).

**Figure 6.**
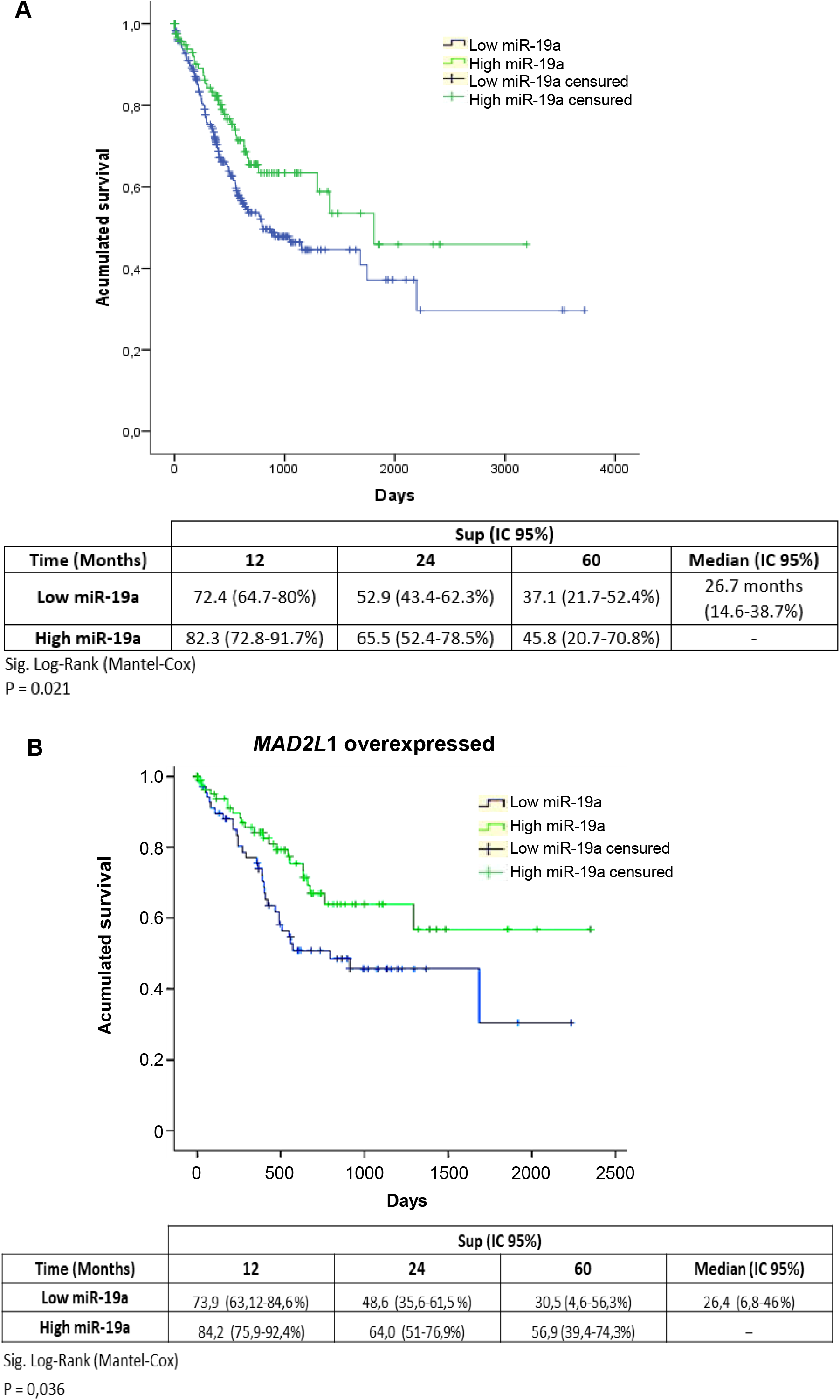
Low miR-19a predicts better prognosis. **A)** Kaplan-Meier analysis with the log-rank test indicated that low miR-19a expression (n = 250, Blue lines), had a significant impact on overall survival (P = 0.021). **B)** Kaplan-Meier analysis of 5-year overall survival curves of patients with high *MAD2L1* and high miR-19a versus high *MAD2L1* and low miR-19a patients of public database TCGA for STAD (stomach adenocarcinoma) mRNAseq illuminahiseq_rnaseqv2-RSEM_genes_normalized (MD5). The statistical significance was evaluated with Mantel-Cox test.

The results suggest that the ratio of *MAD2L1*/miRNA19-a in GC patients correlates with poor clinical outcome, and miR-19a could be considered as a prognostic biomarker.

## Discussion

Accurate MAD2 is crucial to maintain mitosis control and its deregulation increases mitotic errors and affects the recurrence-free survival in GC [8, 9] and other types of cancer [23, 24]. Now, we hypothesize that the over-expression of MAD2 is produced by post-transcriptional modifications through miRNAs and we validate such hypothesis, first with *in silico* analysis, then with *in vitro* functional studies and finally with clinical data, observing that miR-19a is an efficient *MAD2L1* regulator in GC cells.

By *in silico* analysis, we could identify two miRNAs (miR-19a and miR-203) that participate in *MAD2L1* expression. It have been described that *MAD2L1*-3’ UTR region presents binding sequences to different miRNAs, for example: miR-28-5p, miR-493-3p, miR-200c-5p, miR-133b and miR-139a in B-cell lymphomas, breast and ovarian cancer, hepatocellular carcinoma, colorectal and lung adenocarcinoma [22, 25-28]. We decided to include at least one of the previous published miR (miR-139) although our technical approaches did not identify miR-139, to reinforce our data on miR-19a and miR-203 and their physiological role of them in GC [29]. Finally, other miR’s could be involved in GC tumorigenesis with more than one target, for instance miR-30a-3p have been described as *MAD2L1* and APBB2 regulator in lung and gastric cancer respectively[30]. The novelty of our findings is that we have identified additional miRNAs related to *MAD2L1* expression in GC and moreover, we identified its role in the disease in different positions in the *MAD2L1* 3’ UTR (miR-19a at 189bp and miR-203 at 475bp).

In recent years, miRNAs have attracted increasing attention in cancer-related studies, as they effectively regulate several pathological and biological processes, and their molecular targets associated with signaling pathways regulation, gene expression, prognosis, and potential therapeutic approaches for cancers, including GC [31-34]. On these bases, we found two different miRNAs (19a and 203) related to MAD2 expression to be potentially used as a predictive biomarker in GC patients.

The treatment election in GC remained a challenge in spite that several chemotherapeutic regimens have been established [35]. Some of the most used regimens are paclitaxel (PXL) or docetaxel combined with fluorouracil plus cisplatin (PCF) and oxaliplatin combined with other drugs [36-38]. Clinical trials have shown an improvement in response to drug combinations [39]. However, none of these combinations have substantially improved the cure of the disease, and to that end, our group have been working in the designs of metallodrugs, such as I5 and I6 [20]. We have used these metallodrugs in our GC studies because the anti-tumor agents’ sensitivity could also be modulated by miRNAs and/or MAD2 expression. For example,miR-19a plays a critical role in the mechanism of arsenic trioxide treatment in bladder cancer [40]; overexpression of the miR-493-3p conferred cancer cells resistance to PXL [26]; silencing MAD2 enhance the PXL cytotoxicity [8] and miR-200c-5p decreases human hepatocellular carcinoma (HCC) tumorigenesis via suppressing *MAD2L1* [27]

Our data showed a synergy of the platinum derviatives in cell death when we combined the miR-19a and miR-203 expression with G2/M target drugs (I5 and I6). Although these Pt(II) derivatives share an S-phase checkpoint activation, like cisplatin, they also activate the G2/M checkpoint, like PXL. Taken together, the data suggest that GC patients with low/normal levels of MAD2 could be beneficiated by the treatment with mitotic checkpoint activating therapies, opening doors to the discovery of new combination therapies targeting MAD2 in combination with PXL, classical chemotherapeutic agents or new designs.

The role of miRNAs on metastasis is widely reported, for example, the overexpression of miR-320a [28, 41] and miR-335 [42, 43] inhibit breast cancer invasiveness, metastasis and migration *in vitro* and *in vivo*. However, in pancreatic cancer, the overexpression of miR-320a contributes to cancer development, indicating that the same miRNA can even have opposite functions depending on the tumor type [44, 45]. We have observed a reduction in cell mobility (migration & invasion) in GC cells predictably *via* miR-19a/*MAD2L1*. This effect could be potentiated along with other miRNAs targeting *MAD2L1*, like the previously mentioned miR-30a-3p that could decrease the cell proliferation by modulating MAD2 expression[30]. Although miR-203 decreases MAD2 levels, it does not appear to have a specific function in this context, possibly because it may affect other MAD2-independent molecular targets or require alternative pathways [25, 26, 46, 47]. Further approaches, like large-scale and long-term follow-up studies are needed to confirm the significance of miR-203 in GC.

Our analysis of OS using the Kaplan-Meier plot, in a pilot series of patients, have shown that *MAD2L1* overexpression produces a decrease in OS. These results were endorsed by using KM plot.com. We suggest an inverse correlation between miR-19a and *MAD2L1* levels that match with OS, and although we must expand the number of patients to perform further studies, we provide the first evidence supporting the role of miR-19a as a MAD2 regulator and prognostic biomarker in GC targeting *MAD2L1*.

The results found allow us to propose a working model (Fig. 7) in which miRNA-19a could serve as a predictive biomarker in GC. Tumors with a low expression of miR-19a would cause overexpression of MAD2 protein, favoring the capacity of invasion and proliferation of cancer (Fig. 7 bad prognostic). In a scenario in which MAD2 levels were unaltered, it would serve as a biomarker of response to therapy. In a hypothetical situation it could be combined drugs oriented to potentiate the expression of these miRNA and drugs that affect the MC such as PXL or different platinum derivatives (Fig. 7 better pronostic).

**Figure 7.**
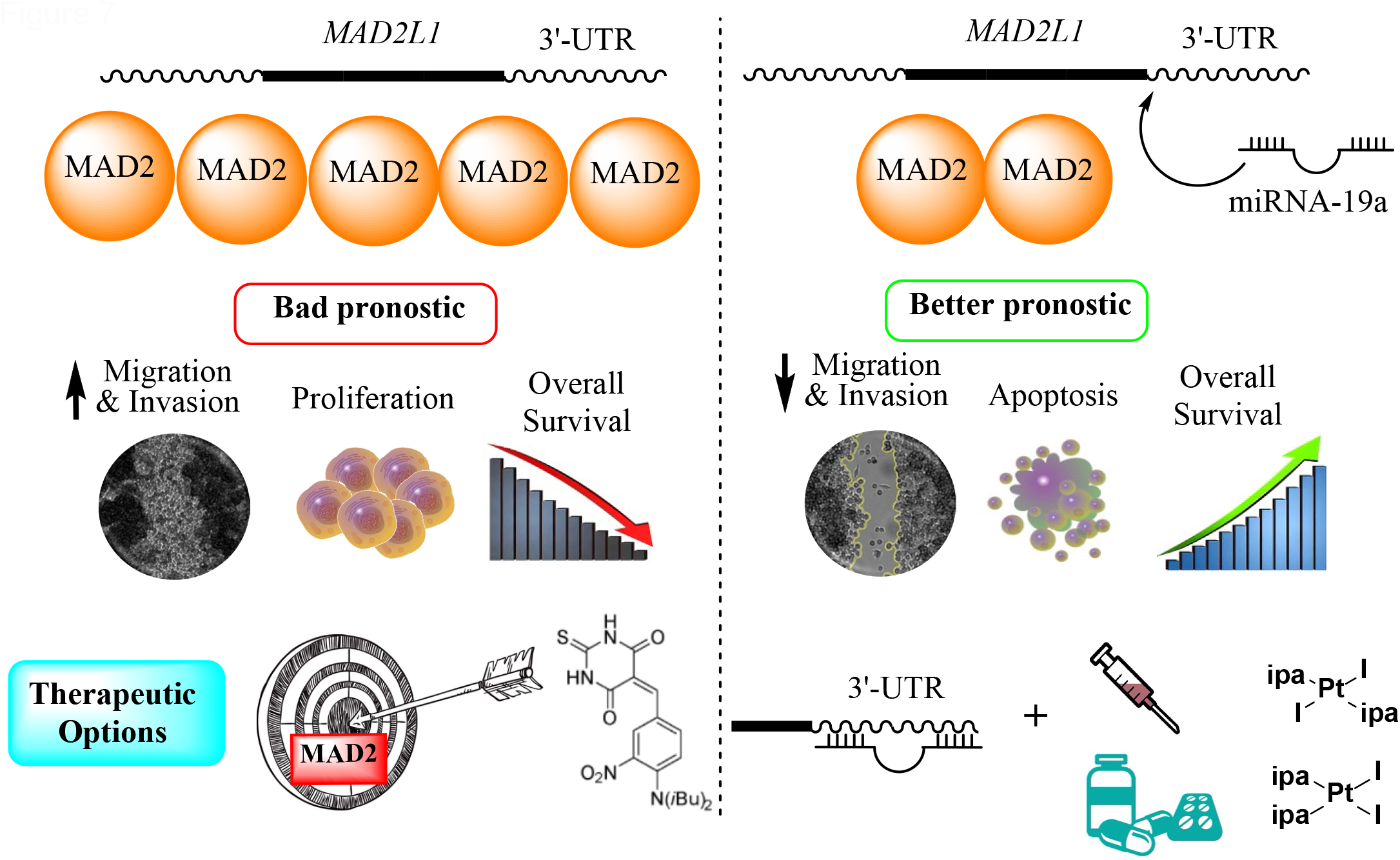
Graphical Summary. Working model of how MAD2 downregulation due to miR-19a binding in the 3’-UTR region enhances OS decreasing proliferation and cell migration and invasion in GC (right panel). The combination with other G2/M drugs like PXL or new platinum derivatives could have a synergistic effect in cell death. The inverse situation is shown on the left side, and the higher expression of MAD2, even being a worst prognostic, could be overcome if it was detected on early stages of the disease applying MAD2-target drugs such as M2I-1 (drafted in the picture).

In summary, the large amounts of data regarding molecular research on GC prognosis and drug response need to be integrated into clinical practice. Overexpression of MAD2 in GC patients is an indicative of poor prognosis and is regulated among other factors by miR-19a and miR-203. Remarkably, modulation of miR19-a decrease migration and invasion and its combination with the platinum drug treatment modulates apoptosis. The translational studies of all these data into clinical practice is the challenge we face in the near future.

## Author Contributions

NPL, JBI, JMH, CC and MP performed experiments. AGQ, RP and ISP devised study designs and wrote the manuscript.

## Institutional Review Board Statement

“The study was conducted according to the guidelines of the Declaration of Helsinki and approved by the Ethics Committee: Comité de ética de investigación de la Universidad Autónoma de Madrid with reference number (approval number) CEI69-1224 (approval date 15/02/2016)).” Approval was waived for this study as Not applicable by Instituto nacional de cancerología Board Statement, reference number INCAN/ CI /817/16 with a not applicable consent (approval date 20/10/2016)

## Acknowledgments

This research was funded by Spanish MINECO grant number PID2019-106220RB100.and RTC2019-007227-1. This work was supported by, and with Fondo de Investigaciones Sanitarias (FIS), Instituto de Salud Carlos III, Spain, grant PI20 0335, (all co-financed through Fondo Europeo de Desarrollo Regional (FEDER) “Una manera de hacer Europa”). NPL was supported by a grant FPU15/04669, funded by Ministerio de Educación, Cultura y Deporte, Spain. JMH FPI-UAM 2021 from Universidad Autónoma de Madrid (Molecular BioSciences PhD programme). We would also to thank the productive scientific discussion from meetings and collaboration from MetDrugs Network RED2018-102471T and COST STRATAGEM CA-17104 and COST NECTAR.

We thank Javier Perez (IIBm Image facility), Monica Belinchón and Lucía Guerrero (IIBm Microscopy facility) for their technical assistance. We are grateful to Dr. Luis del Peso for their generous knowledge support and Dr. Luis Alonso Herrera for Patients RNA samples (INCAN-Mexico).

## Conflicts of Interest

The authors declare no conflict of interest.

## Notes

### Competing Interest Statement

The authors have declared no competing interest.

