## Supplementary figures and images for "Regulatory effects of miRNA-19a on MAD2 expression and tumorigenesis in gastric cancer"

### SM1

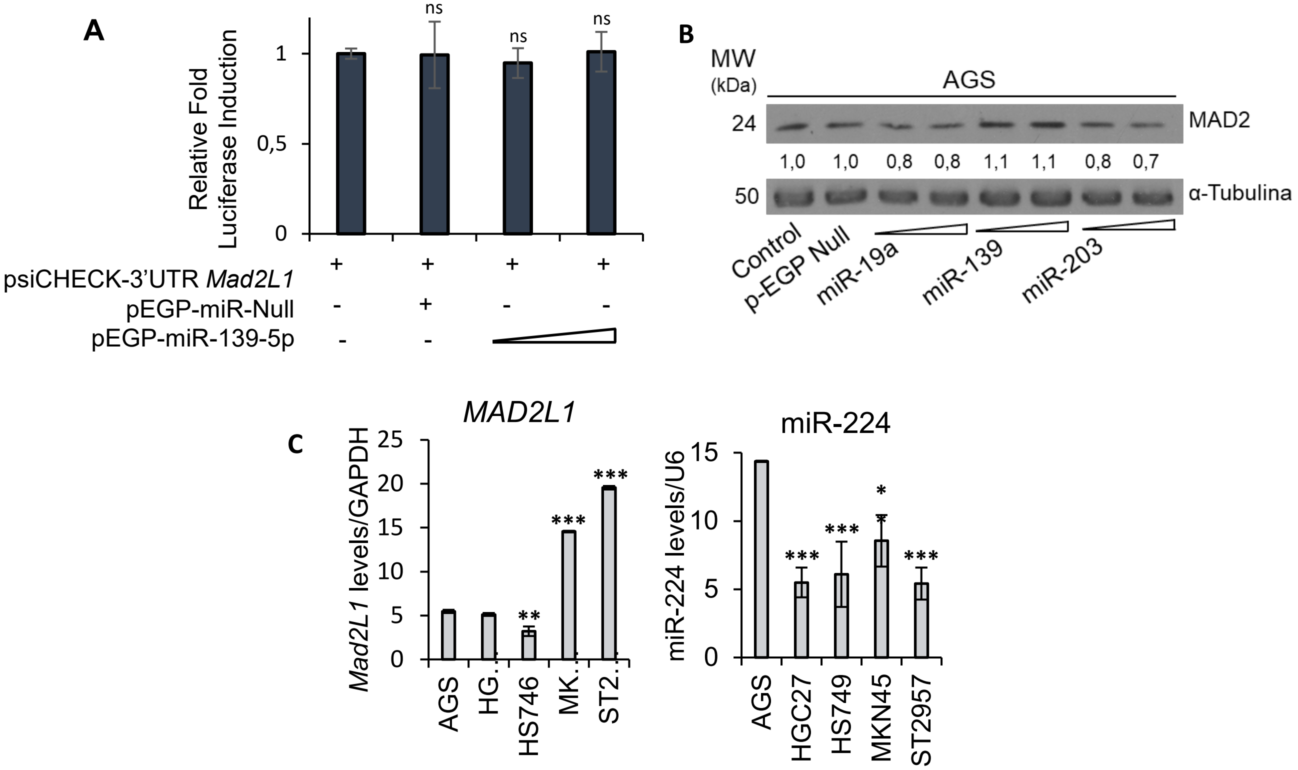
