## Supplementary material for "Regulatory effects of miRNA-19a on MAD2 expression and tumorigenesis in gastric cancer": Tables

| HITS | pvalue | FRV | miRNA ID |
|---|---|---|---|
| 7464 | 5,8757E-07 | 0,00134907 | hsa-miR-1 |
| 10884 | 1,3864E-05 | 0,03183235 | |
| 3914 | 5,2066E-06 | 0,0119543 | hsa-miR-106a |
| 6252 | 2,0465E-05 | 0,04698808 | |
| 5258 | 4,4911E-09 | 1,0312E-05 | hsa-miR-106b |
| 2450 | 5,6566E-09 | 1,2987E-05 | |
| 12522 | 1,2876E-08 | 2,9564E-05 | |
| 13127 | 2,3078E-08 | 5,2987E-05 | |
| 15061 | 2,5391E-10 | 5,8298E-07 | hsa-miR-133a |
| 5279 | 7,9968E-10 | 1,8361E-06 | |
| 13869 | 6,9775E-07 | 0,00160203 | |
| 3787 | 7,4918E-07 | 0,00172012 | |
| 3099 | 3,401E-06 | 0,00780876 | hsa-miR-133b |
| 4117 | 1,7371E-08 | 3,9885E-05 | |
| 8667 | 3,2719E-08 | 7,5123E-05 | |
| 5331 | 1,1264E-08 | 2,5863E-05 | hsa-miR-144 |
| 12158 | 1,2999E-08 | 2,9845E-05 | |
| 4473 | 4,9347E-08 | 0,0001133 | |
| 5552 | 1,8013E-07 | 0,00041359 | |
| 12691 | 2,1691E-10 | 4,9804E-07 | hsa-miR-146a |
| 3827 | 1,2488E-09 | 2,8673E-06 | |
| 7347 | 6,4206E-09 | 1,4742E-05 | |
| 2870 | 1,5715E-08 | 3,6082E-05 | |
| 6092 | 2,5312E-11 | 5,8117E-08 | hsa-miR-148a |
| 4621 | 3,1224E-11 | 7,1689E-08 | |
| 1248 | 1,0279E-10 | 2,36E-07 | |
| 14402 | 1,6822E-10 | 3,8622E-07 | |
| 3869 | 2,0665E-07 | 0,00047448 | hsa-miR-17-5p |
| 354 | 2,3345E-06 | 0,00535993 | |
| 1679 | 2,9284E-06 | 0,00672364 | |
| 15054 | 5,311E-07 | 0,00121941 | hsa-miR-185 |
| 14844 | 7,2434E-06 | 0,01663088 | |
| 1313 | 1,142E-05 | 0,02622021 | |
| 6439 | 9,4062E-09 | 2,1597E-05 | hsa-miR-18a |
| 1980 | 9,9534E-09 | 2,2853E-05 | |
| 4222 | 4,2124E-08 | 9,6716E-05 | |
| 9383 | 1,4136E-07 | 0,00032456 | |
| 13087 | 9,731E-08 | 0,00022342 | hsa-miR-18b |
| 14524 | 1,1231E-07 | 0,00025787 | |
| 11846 | 2,0159E-07 | 0,00046286 | |
| 11909 | 7,8257E-07 | 0,00179679 | |
| 10396 | 6,6152E-07 | 0,00151886 | hsa-miR-195 |
| 10952 | 3,3604E-06 | 0,00771545 | |
| 15719 | 1,2906E-05 | 0,02963225 | |
| 10693 | 1,3694E-06 | 0,0031442 | |
| 11055 | 4,9087E-06 | 0,01127031 | |
| 14346 | 8,2269E-06 | 0,01888893 | hsa-miR-19a |
| 1692 | 1,6807E-05 | 0,03858971 | |
| 2778 | 2,114E-05 | 0,04853705 | |
| 10158 | 2,7728E-05 | 0,063663 | |
| 3525 | 1,8358E-06 | 0,00421498 | hsa-miR-203 |
| 11940 | 2,2472E-06 | 0,00515965 | |
| 10277 | 3,4028E-06 | 0,00781276 | |
| 9857 | 6,6941E-06 | 0,01536957 | |
| 6880 | 2,9458E-08 | 6,7636E-05 | hsa-miR-204 |
| 11597 | 3,383E-07 | 0,00077673 | |
| 6886 | 1,3403E-06 | 0,00307733 | |
| 2688 | 2,0291E-06 | 0,00465887 | |

| | | | |
|---|---|---|---|
| 13208 | 4,4565E-12 | 1,0232E-08 | hsa-mir-25 |
| 8050 | 4,7768E-12 | 1,0968E-08 | |
| 3601 | 5,6546E-12 | 1,2983E-08 | |
| 602 | 2,5691E-08 | 5,8988E-05 | |
| 11819 | 3,0769E-15 | 7,0645E-12 | hsa-mir-375 |
| 4351 | 3,5535E-15 | 8,1588E-12 | |
| 1561 | 7,2315E-15 | 1,6604E-11 | |
| 8444 | 3,7637E-14 | 8,6414E-11 | |
| 3661 | 8,2477E-06 | 0,0189367 | hsa-miR-424 |
| 14424 | 1,2307E-13 | 2,8256E-10 | hsa-mir-451 |
| 13461 | 6,3226E-13 | 1,4517E-09 | |
| 1934 | 1,2929E-12 | 2,9684E-09 | |
| 2468 | 2,1241E-12 | 4,8769E-09 | |
| 10306 | 7,9297E-06 | 0,01820667 | hsa-miR-484 |
| 9411 | 4,4421E-10 | 1,0199E-06 | hsa-miR-486 |
| 15289 | 1,1197E-09 | 2,5709E-06 | |
| 6178 | 2,7071E-07 | 0,00062156 | |
| 15227 | 7,2507E-07 | 0,00166477 | |
| 11913 | 1,4051E-06 | 0,00322601 | hsa-miR-497 |
| 12803 | 1,4954E-05 | 0,03433485 | |
| 6541 | 4,7729E-07 | 0,00109585 | hsa-miR-503 |
| 4282 | 2,4927E-06 | 0,00572325 | |
| 14515 | 6,8836E-06 | 0,01580473 | |
| 8706 | 2,5723E-05 | 0,05906049 | |
| 4873 | 8,6567E-06 | 0,01987584 | hsa-miR-552 |
| 408 | 1,0244E-05 | 0,02352128 | hsa-miR-625 |
| 11771 | 1,4948E-09 | 3,4321E-06 | |
| 10370 | 2,6229E-09 | 6,0222E-06 | |
| 10928 | 1,0059E-10 | 2,3096E-07 | |
| 530 | 8,7002E-06 | 0,01997572 | hsa-miR-650 |
| 1362 | 8,7108E-06 | 0,02000009 | hsa-miR-769-5p |
| 3755 | 5,9602E-07 | 0,00136847 | hsa-miR-92 |
| 2849 | 6,184E-07 | 0,00141985 | |
| 88 | 1,368E-05 | 0,03141025 | |
| 2110 | 1,8854E-06 | 0,00432881 | |
| 6085 | 1,0905E-09 | 2,5038E-06 | hsa-miR-93 |
| 1247 | 3,0148E-09 | 6,922E-06 | |
| 10677 | 4,4865E-08 | 0,00010301 | |
| 13858 | 4,5796E-06 | 0,01051486 | |

| |
|---|
| hsa-miR-1 |
| hsa-miR-106a |
| hsa-miR-106b |
| hsa-miR-133a |
| hsa-miR-133b |
| hsa-miR-144 |
| hsa-miR-146a |
| hsa-miR-148a |
| hsa-miR-17-5p |
| hsa-miR-185 |
| hsa-miR-18a |
| hsa-miR-18b |
| hsa-miR-195 |
| hsa-miR-19a |
| hsa-miR-203 |
| hsa-miR-204 |
| hsa-mir-224 |
| hsa-mir-25 |
| hsa-mir-375 |
| hsa-miR-424 |
| hsa-mir-451 |
| hsa-miR-484 |
| hsa-miR-486 |
| hsa-miR-497 |
| hsa-miR-503 |
| hsa-miR-625 |
| hsa-miR-769-5p |
| hsa-miR-92 |
| hsa-miR-93 |

| miRNA ID | Target Scan/miRBase |
|---|---|
| hsa-miR-203 | MAD2L1 |
| hsa-miR-19a | |
| hsa-miR-625 | |
| hsa-miR-148a | |
| hsa-mir-224 | |
| hsa-miR-1 | OT* |
| hsa-miR-106a | |
| hsa-miR-106b | |
| hsa-miR-133a | |
| hsa-miR-133b | |
| hsa-miR-144 | |
| hsa-miR-146a | |
| hsa-miR-17-5p | |
| hsa-miR-185 | |
| hsa-miR-18a | |
| hsa-miR-18b | |
| hsa-miR-195 | |
| hsa-miR-204 | |
| hsa-mir-25 | |
| hsa-mir-375 | |
| hsa-miR-424 | |
| hsa-mir-451 | |
| hsa-miR-484 | |
| hsa-miR-486 | |
| hsa-miR-497 | |
| hsa-miR-503 | |
| hsa-miR-769-5p | |
| hsa-miR-92 | |
| hsa-miR-93 | |

* Other targets
